# HTS-IBIS: fast and accurate inference of binding site motifs from HT-SELEX data

**DOI:** 10.1101/022277

**Authors:** Yaron Orenstein, Ron Shamir

**Affiliations:** Blavatnik school of computer science, Tel-Aviv University, Israel

## Abstract

**Summary:** Recent technological advancements enable measuring the binding of a transcription factor to thousands of DNA sequences, in order to infer its binding preferences. High-throughput-SELEX measures protein-DNA binding by deep sequencing over several cycles of enrichment. We devised a new algorithm called HTS-IBIS for the inference task. HTS-IBIS corrects for technological biases, selects the cycle and k, and builds a motif starting from a consensus k-mer in that cycle. In large scale tests, HTS-IBIS outperformed the extant automatic algorithm for the motif finding task on both *in vitro* and *in vivo* binding prediction.

**Availability:** HTS-IBIS is available on acgt.cs.tau.ac.il/HTS-IBIS.

## 1 INTRODUCTION

Recent technological advancements provide high-resolution measurements on the binding preference of a specific transcription factor (TF) to thousands of double-stranded DNA oligos. Protein binding microarrays (PBMs) measure *in vitro* binding to thousands of microarray probes (Berger *et al.* 2006). ChIP-seq technology sequences *in vivo* bound genomic sequences (Barski and Zhao 2009). The most recent of these technologies, high-throughput-SELEX (HTS), utilizes deep sequencing to measure TF binding preference through several cycles of enrichment (Zhao *et al.* 2009; Jolma *et al.* 2010; Slattery *et al.* 2011). Jolma *et al.* (2010) describe an algorithm, which we call *Toivonen’s algorithm*, for this task. That algorithm was later used in (Jolma *et al.* 2013) to infer binding site motifs in 547 experiments. However, for a fraction of the reported motifs the starting solution (seed) was manually picked (E. Ukkonen, private communication).

Here we describe a new fully automatic algorithm called HTS-IBIS (HTS Inference of BInding Sites) for inferring a binding model from HTS data, while addressing systematic biases, such as over-concentration and sequence biases. We compare HTS-IBIS to Toivonen’s algorithm and to the models published in (Jolma *et al.* 2013).

## 2 APPROACH

First, we describe our strategy to overcome systematic biases in HTS data. The k-mer distribution in oligos of the initial cycle is near-random. As cycles advance it becomes more skewed towards the binding site, with a risk of eventual over-concentration on one or few k-mers. To select the best cycle we compute KL-divergence score of the 100·4^k-6^ most frequent k-mers in each cycle for k=6-8. The first cycle with a score > 0.1 (or the last, if all cycles have a score ≤ 0.1) is chosen. Additionally, in (Orenstein and Shamir 2014) we identified systematic biases in k-mer frequencies in HTS data. We address such biases here using the observation that they are strand-specific. To score k-mers, we use counts and not ratios between counts in consecutive cycles, since we observed that the former are more correlated to PBM binding intensities. For every k-mer *S* with count *c*(*S*) and its reverse complement 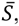 we replace *c*(*S*) by min 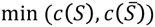 if either 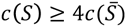 or *S* has k-2 identical nucleotides or more.

HTS-IBIS works in stages: (i) The cycle to work on is found as described above. (ii) k is chosen in the selected cycle as in (Slattery *et al.* 2011). (iii) The k-mer counts are corrected as above and the most frequent k-mer (*consensus* or *seed*) is found. (iv) PWM generation is akin to our RAP algorithm for PBM data (Orenstein *et al.* 2013): a set of 20·4^k-4^ top scoring k-mers is aligned to the seed and nucleotide probabilities in each column are calculated based on their relative scores in the aligned position. (v) The PWM is extended to the sides, and then uninformative side positions are trimmed. An efficient implementation of the software runs in ~5 seconds. See Supplementary Information for details.

## 3 RESULTS

We first assessed our seed finding process. We calculated the success rate in finding a seed that fits (a) the top-ranking 8-mer in a PBM experiment on the same TF, and (b) one of the seeds published by Jolma *et al.* (2013) for the same TF. In both cases, our seed finding process (step (i)-(iii) above) achieved increased success rate compared to selecting the most frequent 8-mer (Figure 1A and Supplementary Table 1).

**Fig 1.**
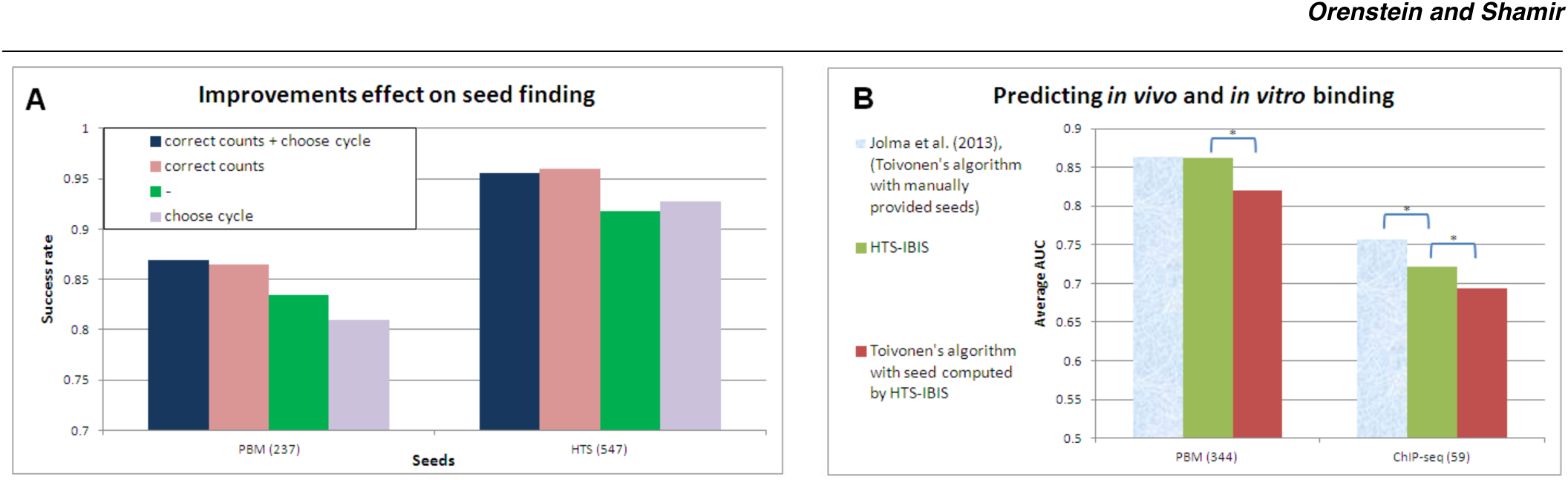
Success rate in seed finding and quality of binding prediction. A) Accuracy in seed finding, with and without bias correction and cycle selection. Success is defined as finding a seed similar (i.e. different in ≤2 positions, and with ≤2 positions offset) to the top-ranking 8-mer in a PBM experiment on the same protein (tested on 237 paired PBM-HTS experiments) or to the seeds published by Jolma *et al.* (tested on 547 HTS experiments). B) Binding prediction quality of HT-IBIS, Toivonen’s algorithm, and models published in Jolma *et al.* (2013), which used some manual inputs. Performance was evaluated by AUC for predicting *in vitro* and *in vivo* binding (tested on 344 paired PBM-HTS and 59 paired ChIP-seq-HTS experiments).

Second, we compared the PWMs produced by HTS-IBIS to two others: (1) Toivonen’s algorithm (Jolma *et al.* 2010), which generates a model given a seed. Since we could not reproduce the seed finding process in (Jolma *et al.* 2010), we ran that algorithm providing the consensus obtained in step (iii) of HTS-IBIS as a seed. (2) The models published in (Jolma *et al.* 2013). To achieve these results, the authors used Toivonen’s algorithm, but occasionally with manual intervention to determine the seed. We used these models to predict PBM binding (Robasky and Bulyk 2011) and ChIP-seq binding (Landt *et al.* 2012). The algorithm of (Slattery *et al.* 2011) was excluded from the comparison since it uses an ‘all k-mers’ model, which has far more parameters than a PWM model. The results are summarized in Figure 1B and Supplementary Table 2. For predicting PBM binding, HTS-IBIS is on par with the published models, while significantly better than Toivonen’s algorithm. As for *in vivo* binding prediction, the published models perform significantly better than HTS-IBIS, which outperforms Toivonen’s algorithm.

## 4 DISCUSSION

HTS technology produces measurements of *in vitro* binding on an unprecedented scale, but suffers from systematic biases. Over-concentration may occur at high cycles, where only high-affinity k-mers may be captured. Biases in oligo generation, sequencing and PCR may explain the observed abundance of A-rich and C-rich k-mers. We developed ways to address these biases. We use the KL-divergence score to choose a cycle that is not too specific, and we correct inflated k-mer counts by comparing them to their reverse complements and by down-scaling degenerate ‘sticky’ k-mers.

We developed the HTS-IBIS algorithm to infer a binding model from the data, and tested it extensively on both *in vivo* and *in vitro* binding prediction. The models published in (Jolma *et al.* 2013) performed best in both tasks. However, since some of these models used external information on the seed, the comparison is improper: one cannot make a meaningful comparison between algorithms that work automatically and algorithms that receive additional external information. Toivonen’s algorithm, run on the same set of seeds, performed worse than HTS-IBIS in both tasks. We believe this is due to the ‘strictness’ of models produced by Toivonen’s algorithm, which uses only k-mers at Hamming distance at most 1 from the consensus.

The rich available HTS data provide a great opportunity to advance our understanding of TF–DNA binding. Improved models combining additional considerations, e.g. biomechanical and structural properties, may be developed in the future.

## ACKNOWLEDGEMENTS

We are grateful to J. Toivonen for providing us with his software. We thank E. Ukkonen for discussions of the algorithm.

### Funding

Israel Science Foundation (ISF) [317/13] (to R.S.); Edmond J. Safra Center for Bioinformatics at Tel Aviv University, the Dan David Foundation, and the Israeli Center for Research Excellence (I-CORE), Gene Regulation in Complex Human Disease, center 41/11 (to Y.O.).

